# Measuring Capillary Flow Dynamics using Interlaced Two-Photon Volumetric Scanning

**DOI:** 10.1101/2022.09.12.507580

**Authors:** John T Giblin, Seong-Wook Park, John Jiang, Kıvılcım Kılıç, Sreekanth Kura, David A. Boas, Ichun A. Chen

## Abstract

Two photon microscopy and optical coherence tomography (OCT) are two standard methods for measuring flow speeds of red blood cells in microvessels, particularly in animal models. However, traditional two photon microscopy lacks the depth of field to adequately capture the full volumetric complexity of the cerebral microvasculature and OCT lacks the specificity offered by fluorescent labeling. In addition, the traditional raster scanning technique utilized in both modalities requires a balance of image frame rate and field of view, which severely limits the study of RBC velocities in the microvascular network. Here, we overcome this by using a custom two photon system with an axicon based Bessel beam to obtain volumetric images of the microvascular network with fluorescent specificity. We combine this with a novel scan pattern that generates pairs of frames with short time delay sufficient for tracking red blood cell flow in capillaries. We track flow speed in 10 or more capillaries simultaneously at 1 Hz in a 237 μm x 237μm x 120 μm volume and quantify both spatial and temporal variability in speed. We also demonstrate the ability to track flow speed changes around stalls in capillary flow.

## Introduction

Capillary blood flow is essential for supplying the brain with oxygen and other nutrients needed to support the high metabolic demand required for normal brain function^1^. Numerous studies have shown an association between reduced cerebral blood flow and cognitive decline^2,3^, including changes in microvasculature structure and flow^4–6^. More recently, transient disruptions in flow at the single capillary level were identified as a potential mechanism that contributes to cerebral blood flow changes driving neurological deficits^7,8^. In addition to increasing vascular resistance, capillary flow models suggest that increased heterogeneity of capillary transit times, which stalls would induce, reduces the efficiency of the microvascular network^9–11^. However, experimental observations of both spatial and temporal capillary flow speed variability across a large volumetric field of view (FOV) with numerous capillaries in the microvascular network is missing. This is because traditional multiphoton microscopy used to measure red blood flow speeds in capillaries^12,13^ uses line scans along the axis of a capillary to observe the passage of RBCs through a single capillary. As a result, many of the capillary flow speeds reported are from measurements of flow speeds where a capillary is monitored for as little as 30 seconds, and in a sequential manner. Though capillary flow speed measurements have been extended to measure RBCs in multiple capillaries simultaneously by employing different scanning sequences^14–16^ the short axial point spread function limits the depth of field; thus only a limited number capillaries can be measured at a single depth in a given time.

Recently, two photon imaging with axially extended Bessel-foci has been shown to greatly increase the volume rate of imaging sparsely labeled samples by allowing for the capture of fluorescent signal in a 3D volume from a single 2D scan. Coupled with a resonant galvo scanner, Fan et al.^17^ showed video rate tracking of RBCs as they flow single file along the capillaries. But tracking speeds of up to 2 mm/s require imaging rates of 100 Hz in a resonant galvo-galvo raster scanning system. At these speeds, the imaging field of view is sacrificed, high excitation powers are required, and averaging to reduce noise will negate the benefits of using resonant galvos.

Another modality to visualize microvascular angiograms is with OCT, which provides volumetrically resolved angiograms from a 2D scan using the endogenous contrast that arises from moving RBCs^18,19^. Doppler^20,21^ and speckle decorrelation^22–24^ OCT imaging have been used to quantify RBC speeds within capillaries but have imaging rates of 10 sec per frame or more^21,25^. This is not suitable for imaging fast flow speed changes within capillaries such as changes due to RBC stalls, which occur on the order of a few seconds or less^26^. Furthermore, OCT lacks the specificity provided by fluorescent labeling that permits identification of the roles of different types of blood cells (erythrocytes, leukocytes, and thrombocytes)^7^ and vessel wall components (e.g. pericytes) in dynamic flow changes^4,27–30^ and disruptions^27,30^.

Our work overcomes these challenges by integrating a novel scanning pattern called “stagger scan” achieved with traditional galvo scanners in an extended Bessel-foci two photon imaging system. Using the stagger scan method we track RBC movement and developed two techniques, Frame Pair (F-Pair) and Frame Tilt (F-Tilt), to measure RBC velocities with average speeds ranging from 0 mm/s to 1.5 mm/s in capillaries across a 237 μm by 237 μm by 120 μm volume. The measured speeds with our method were validated against the traditional line scanning method^12^ for single capillaries. With our approach, RBC speeds across a capillary network were extracted to quantify the spatial and temporal variability in flow speeds, including the ability to capture changes in flow speed due to RBC stalling events both within the stalling and adjacent capillaries. The extracted capillary flow dynamics and distributions matches well with previously published reports^5,11^ while the observed point prevalence in the stall statistics were higher than those reported in^8,26^ due to the increased temporal resolution of our system. Lastly, we demonstrate that the imaging depth is background limited at roughly 400um deep.

## Methods

### Animal Preparation

All animal procedures were approved by the Boston University Institutional Animal Care and Use Committee and were conducted following the Guide for the Care and Use of Laboratory Animals. Following procedures described previously^31^, animals were anesthetized with isoflurane (3% induction, 1-1.5% maintenance) and placed in a stereotaxic frame (Kopf Instruments). Depth of anesthesia was verified and animals were monitored throughout surgery via respiratory rate and toe pinch. Body temperature was maintained at 37 °C with a heating blanket. The craniotomy was performed over one or both hemispheres for optical access to the brain and the dura left intact to minimize damage to vasculature. Exposure was covered with modified Crystal Skull cover glass (LabMaker) and sealed with dental acrylic. For head fixation during imaging, a custom ring-shaped titanium head bar was secured to the remaining cranial bone using low viscosity cyanoacrylate adhesive (Loctite 401) and dental acrylic. Post craniotomy, animals were given a minimum of one week to recover before being acclimated to head fixation. A minimum of 3 weeks was given to allow for further reduction of inflammation after surgery before the animal is imaged.

### In-vivo Imaging

To reduce stress and motion artifacts during imaging, mice were acclimated to head fixation over the course of 7-10 days. During training sessions, animals were secured in a training head fixation jig, fed sweetened condensed milk every 10-15 minutes, and observed for comfort. If the animal exhibited significant discomfort, they were removed from the fixture and the training session was ended. Daily training continued until they were able to comfortably remain in the fixture for 90 minutes.

Blood vessels were labeled with intravenous injection of 50 μL of 150 kDa Fluorescein-Dextran (5% in PBS) ~5-10 minutes prior to imaging. For imaging deeper into the brain (Figure 8), 150 kDa Texas Red-Dextran was used. When imaging, animals were head fixed in a custom designed head fixture with tip, tilt and elevation control adjustment head, and adjusted such that the cranial window orientation is perpendicular to the imaging axis of the microscope. During an imaging session, a treat of sweetened condensed milk was offered to the animal every 30 minute intervals, and immersion fluid was checked and supplemented when necessary.

### Two Photon Microscope and Bessel Foci Generator

Measurements were performed with a custom designed, galvo-galvo, two-photon microscope with an axicon based Bessel foci-generator. The 2P excitation source is from a Spectra-Physics Insight 2 femtosecond laser coupled with an electro-optic modulator (Conoptics 350-105-20) for fast flyback blanking and overall excitation power control. 2P fluorescence from the labeled blood plasma was excited using 920 nm.

Following^32–34^, an axicon (Eskima Optics, Model 131-1279) and a Thorlabs achromat (FL= 250mm) is used to generate the annular ring, which is then imaged onto the first beam steering galvanometer inside the microscope using a Thorlabs achromat (FL= 100 mm) and a Sill Optics lens assembly (S4LFT0089/094) in a 4f configuration. A Thorlabs BE02-05-B beam expander is used to tune the excitation beam diameter before the axicon, which allows us to tune the length of the extended foci from 50 μm to 500 μm without changing optics (see Fig. 1a)

**Figure 1.**
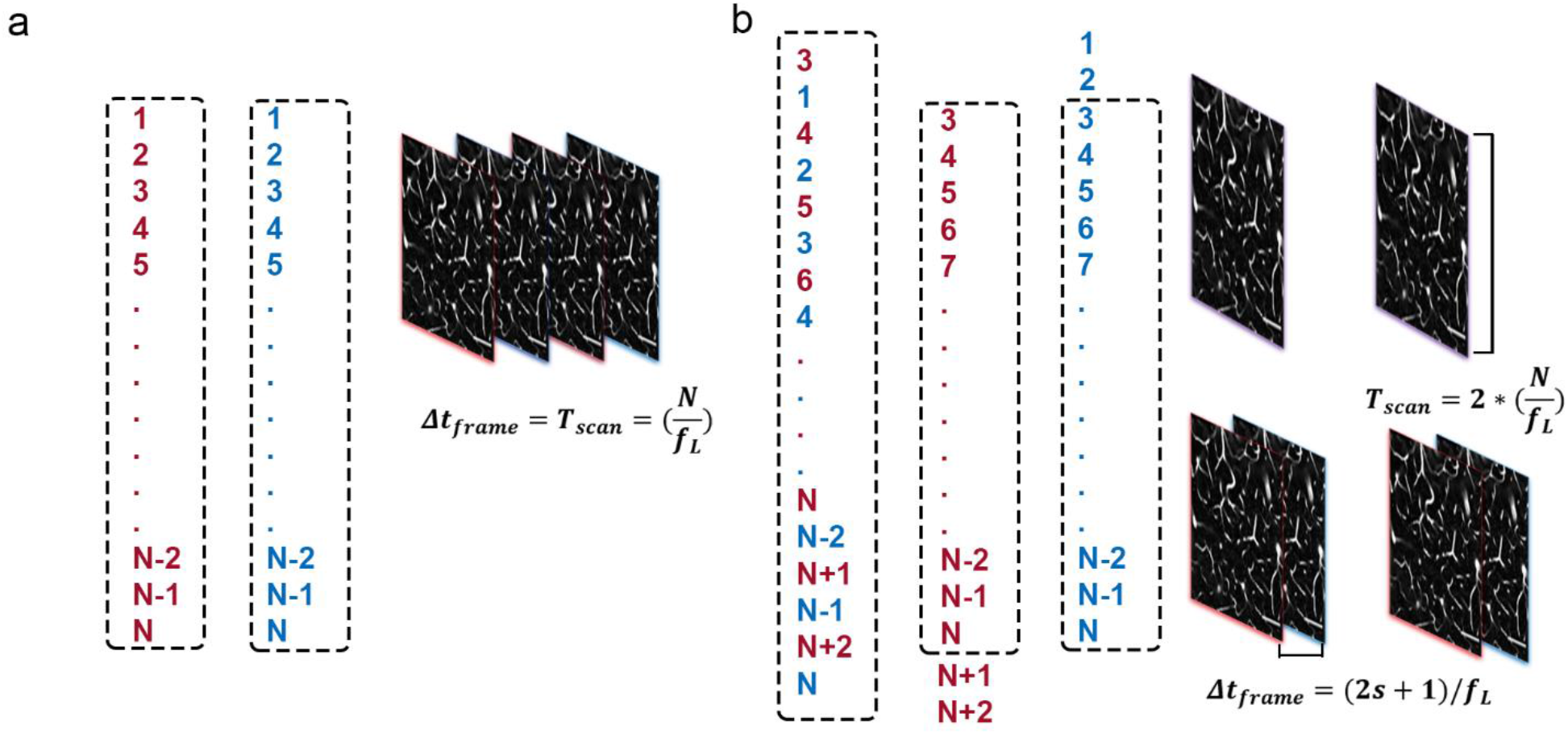
**(a)** Scan sequence for a typical raster scan, all lines are scanned sequentially before returning to the first line and scanning another image **(b)** Scan sequence for a stagger scan. 2 images are constructed simultaneously, with the line jump *s* controlling the number of lines jumped when alternating between the 2. In this case, the time between frames is dramatically reduced.

The microscope consists of two decoupled galvanometers (Thorlabs GVS001) imaged onto each other using two Sill Optics lens assembly (S4LFT0089/094) in a 4f configuration. They are then imaged to the back pupil plane of the objective (Olympus 25x, 1.0NA) using a Sill Optics lens assembly (S4LFT0089/094) and a custom designed Plossl tube lens comprised of two Thorlabs achromates (AC508-500-AB). The objective is mounted on a 1mm, fast piezo Z translation stage for axial scanning (nPoint nPFocus 1000).

For detection of the 2P fluorescent emission, a long-pass primary excitation/emission dichroic (SEMROCK DI03-R785) is placed between the tube lens and objective to reflect the fluorescently emitted light into the detection path optics. For detection optics, two groups of two plano-convex lenses (first group: 2x LA1979-A; second group: 2x LA1131-A) are used to collect the fluorescence onto a red sensitive, cooled PMT (Hamamatsu H7422-50) with an emission filter (SEMROCK FF01-607/70). A secondary dichroic (SEMROCK FF555-DI03) places between optical elements of the first group of plano-convex lenses enables us to add a second detection pathway. In the second pathway, a single plano-convex lens, and pair of plano-convex lenses, collect fluorescent photons onto a second cooled PMT (Hamamatsu H74422-40) with an emission filter (SEMROCK FF01-502/60).

The microscope is controlled using NI industrialized PXIe with an embedded controller (NI PXIe-8880) chassis running ScanImage software^35^. Galvo, pockel cell, and piezo control voltages are generated by ScanImage and sent to their respective control units through a data acquisition card (NI PXIe-6358) with a BNC break out connector box (NI BNC-2110). The photocurrent from the two PMT’s are converted to voltage using Femto DHPCA-100 trans-impedance amplifiers. These voltages, along with feedback position voltages from the galvos are recorded by a second data acquisition card (NI PXIe-6358) with a BNC break out connector (NI BNC-2110).

### Stagger Scanning, Paired Frame Rate, and Measurable Velocities

In conventional unidirectional raster scanned imaging, each line in the image is scanned sequentially until a full frame is generated (Figure 1a). The next frame then begins again at the first line and this sequence is iterated to produce a time series of images. The time delay between two frames is a product of the line rate and the number of lines per image. In a stagger scanned image (Figure 1b), pairs of lines are acquired sequentially with each pair having a fixed line offset, or line jump, that is equal to an integer number of lines. In this manner, two images of the same field of view are constructed simultaneously rather than sequentially. For example, the first pair of lines scanned in the image would represent lines 3 and 1 in image frame (Figure 1b). The next pair of lines in the image would represent lines 4 and 2 in a normal raster scanned image, and so on until all of lines in the scanned image are scanned twice before a single stagger scan frame is complete. So, in a single stagger scanned image, there are two interlaced frames with a pairwise frame delay (PFD) of

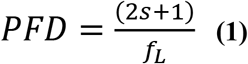

where *f_L_* is the line acquisition rate including flyback overhead in hertz, and s is the line jump. Since *s* is a small number, a PFD on the order of milliseconds can be created at typical raster scanning rates. The total scan period for one stagger scan image becomes

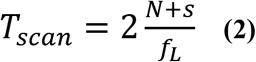

which is roughly equal to the time required for two raster scanned images, but with significantly less time between the two frames. This short time between the pair of frames rate enables tracking of fast-moving RBC’s travelling within capillaries.

To demonstrate the utility of RBC tracking with stagger scanning, we compare the theoretical maximum RBC velocities that can be measured along a 50 μm length of capillary between traditional raster scanning vs stagger scanning for a 256×256-resolution image (Supplemental Figure 1). In the ideal scenario, a single RBC flows along a visible length of capillary between two full image frames and the maximum measurable speed is determined by the length of the capillary and time between the two frames. Assuming a galvo line rate is at 900 Hz, a traditional uni-directional raster has a maximum measurable RBC velocity of ~0.17 mm/s, while in the same scenario, the theoretical maximum RBC velocity with a line jump (*s*) of 2 is roughly 9 mm/s, which is fast enough to resolve any capillary RBC velocities.

Another advantage of stagger scanning with traditional galvos over using resonant scanning galvos for tracking fast dynamics is our ability to control the pixel dwell times when acquiring the image. With a longer pixel dwell time, more fluorescence is generated and translates to a higher contrast between the fluorescent plasma and RBCs. This is particularly important when using an extended axial focus, where excitation power is distributed over a larger volume when compared with traditional Gaussian foci; as well as imaging deep into scattering tissue.

Figure 2a shows an illustration of our axicon based Bessel beam generator module used in our custom built 2P microscope. The elongated axial profile of a 1μm bead is shown and intensity profile is plotted in the graph showing an axial full width half maximum of 100 μm. Comparing the 2P angiograms captured with the extended axial focus with a conventional Gaussian focus (Figure 2b), angiograms generated with the elongated foci capture significantly more capillaries in a single scan. Using a staggered scan, the deinterlaced images create a time series of frame pairs with milliseconds paired frame delay (Figure 2c) that is tunable by changing the line rate of the acquisition, *f_L_*, and the line jump, *s*, between repeated line scans (Equation 1). Within each frame pair, the short time delay allows us to observe RBC shadows as they flow along the length of the capillary (Figure 3a) and estimate RBC flow velocities in multiple capillaries using the configured paired frame delay (PFD) and the measured distances the RBCs travelled along the capillary. We call this method of velocity estimation F-Pair.

**Figure 2.**
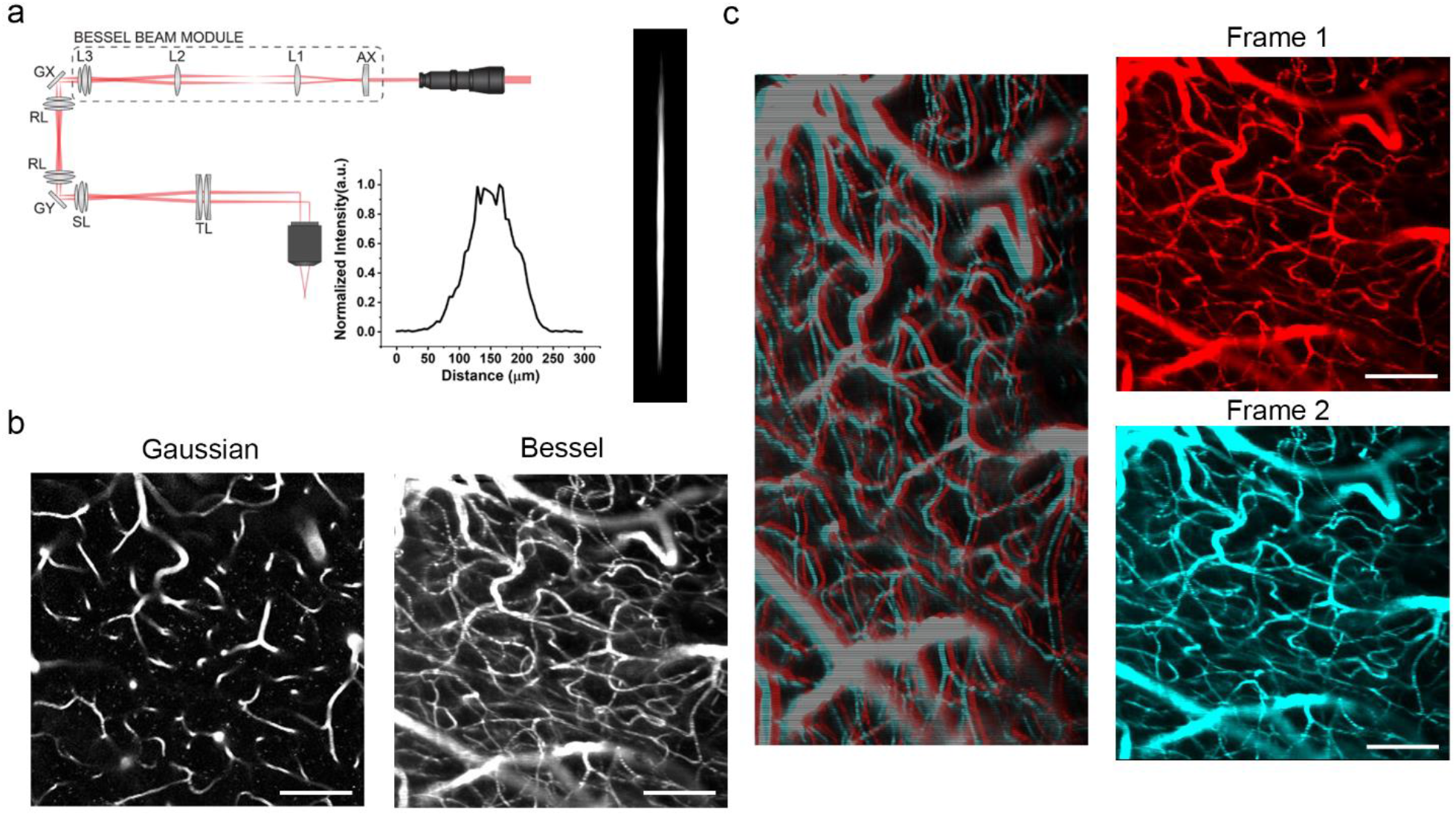
**(a)** Schematic of the Bessel beam two-photon microscope. The axicon (AX) and lens L1 are used to generate the annular illumination required to generate a Bessel like focus, which is imaged onto a plane conjugate to the back pupil plane of the objective of a conventional two-photon microscope using lenses L2 and L3. The Bessel focus creates an axially extended focus shown on the right with the fluorescence intensity trace plotted below showing a full width half maximum of 100 μm **(b)** Comparison of the same ROI imaged with a Gaussian (left) and Bessel (right) focus. The Bessel focus allows for imaging a 3D volume during a single 2D raster scan, visualizing significantly more capillaries in the same field of view (scale bar: 100 μm). **(c)** The staggered scan pattern creates an image containing two interlaced images (left, line phase offset added to make 2 images more clear). This can be separated into a pair of frames (right) generated with a time delay of a few line periods. 512×512 pixels with a paired frame delay (PFD) 12.24 ms. Scale bar 100 μm

**Figure 3.**
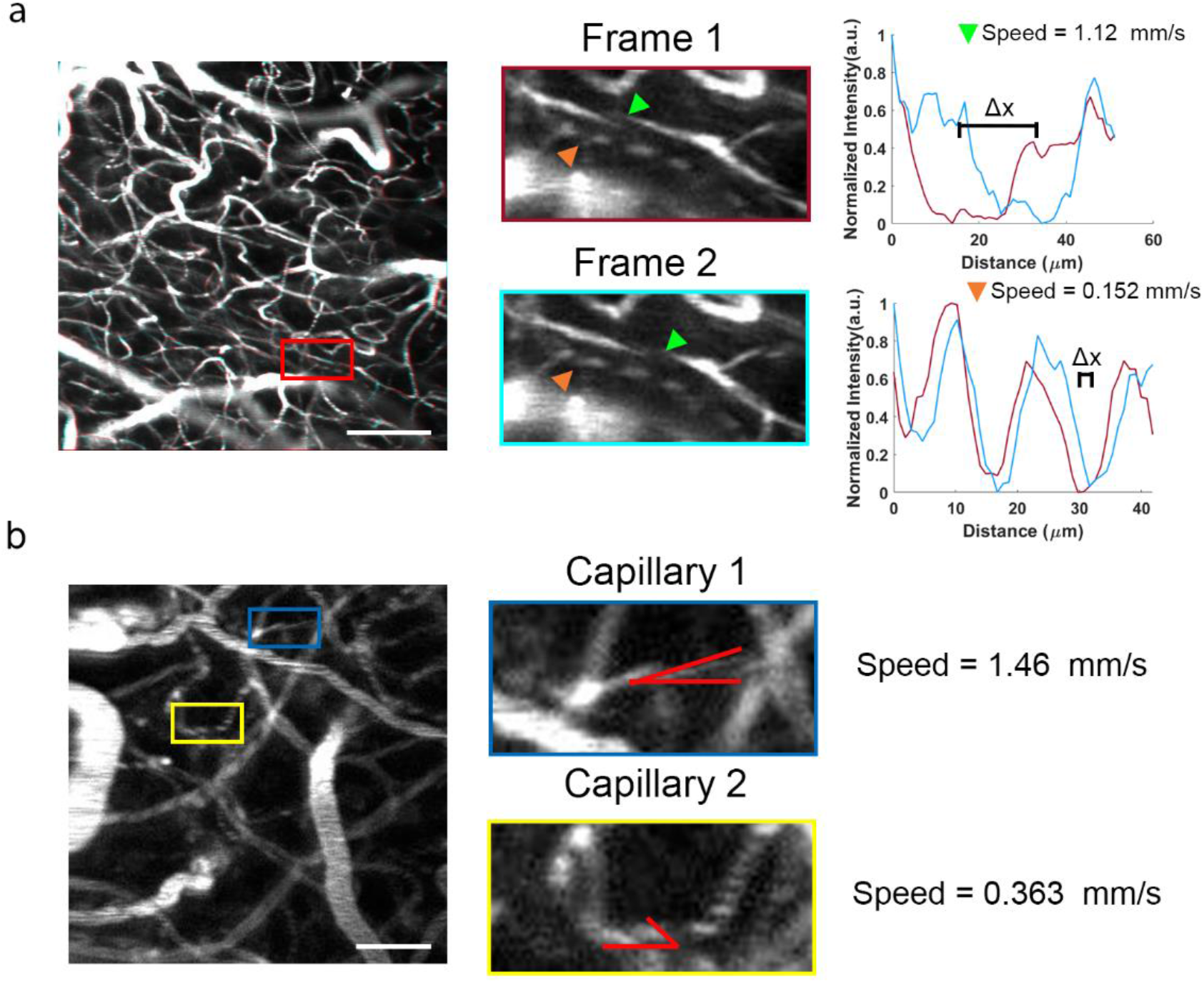
**(a)** The short temporal delay between frame pairs allows for the tracking of RBCs as they flow along a capillary. In fast flowing capillaries (green arrow), RBCs flow a much large distance between frame pairs compared to slower moving capillaries (orange arrow). Taking a centerline profile, the distance Δx the RBC flows can be used to calculated speed using Equation 3. Scale Bar 100 μm **(b)** Comparison of capillaries with different amounts of RBC slewing during image scanning. In the yellow ROI the RBCs were flowing left to right, while in the blue ROI the flow was right to left, as determined by the direction of the slant. By measuring the angle of the slew relative to the capillary axis, the speed can be estimated using Equation 6. Scale bar 50 μm

In addition, the apparent shape of an RBC in a given capillary appears slew due to the RBC flow velocity and the slow axis raster scan velocity (Figure 3b). Faster RBC velocities create a steeper slewed shape (Figure 3b), while slower velocities create a shallower slew shaped. By using this angle of the slew and the known slow axis line rate, an additional method called F-Tilt can be used to estimate flow speed to track RBC flow as previously described in^14–16^.

### RBC velocity measurements using F-Pair and F-Tilt

From the image pair that is deinterlaced from a stagger scan image, there are two methods for estimating RBC speeds. In the F-Pair method, a capillary with an RBC shadow that is visible in both images of the frame pair is identified. The pixel position of the RBCs along the length of the capillary is defined as the center of the RBC shadow in the capillary (Figure 3a). From the pixel positions in each frame pair, the velocity of the RBC along the capillary is simply

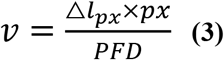

Where Δ*l_px_* is the pixel shift between the two intensity traces in pixels, *px* is the pixel step size in μm/pixel, and PFD is the paired frame delay in seconds. For F-pair to produce reliable speed measurements, RBCs must be uniquely identifiable, otherwise aliasing of the RBC shadows will produce erroneous results.

In the F-tilt method, RBC movement in the capillary will result in an image where the shadow appears skewed in raster scanned systems. Like the image distortion seen with a rolling camera shutter^36^, the faster an RBC is moving through a capillary that is parallel to the fast axes of the scanning, the more the RBC shape is tilted/shifted in the image^14–16^. For a capillary with flow aligned perfectly with the fast axis of scanning, the RBC speed is estimated from

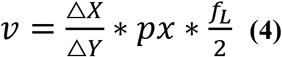

Where Δ *X* is the distance the RBC moved along the length of the capillary in pixels, Δ *Y* is the number of lines it took to scan over the capillary width, and *f_L_* is fast axis line rate in lines/second. When the capillary’s axis does not align perfectly with the imaging fast axis, equation 4 must be modified to account for the orientation of the capillary and the imaging fast axis

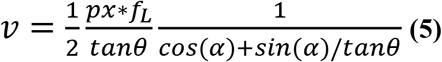

where α is the angle between the capillary and the fast axis accounts for the capillary orientation differences and *θ* is the angle between the RBC shadow and the capillary axis.

According to this equation, F-tilt velocity estimates will have large uncertainties when the angle between the capillary and the fast axis is large. To minimize this error, F-tilt measurements for estimating RBC velocity should be applied to a pair of orthogonally oriented and sequentially acquired stagger scan images.

### Staggered Scanning and Image Reconstruction

The staggered scanning pattern is generated by a custom stimulus function written in Matlab for ScanImage^35^, which computes the galvo control voltages based on user specified parameters defined in ScanImage. The parameters are the total number of lines in one de-interlaced image, the line period (excluding flyback overhead), and the number of lines jumped. For all our experiments, a line period of 512 ms, line jump of 2, total line number of 256, and field of view size of 237.88 μm was used to generate the stagger scan image pairs. This translates to a paired frame delay (PFD) of 5.56 ms for the pairs of 256×256 pixels images.

The acquired data is stored by ScanImage as a raw data file, which is imported into MATLAB using the ScanImage Matlab API. Once the data has been imported into Matlab, a custom script reshapes and deinterlaces the saved data into a time series of frame pairs that is saved as an image TIFF image stack for use in RBC velocity analysis.

### F-Pair Analysis

From the processed TIFF image stack, capillaries of interest are identified and the contrast between the RBC shadow and the fluorescently labeled plasma is visually evaluated. Capillaries with good contrast are then manually annotated with a line ROI along the middle of the capillary using the ROI line tool in ImageJ and the intensity is then extracted for each pair of images in the time series using a custom MATLAB script. The RBC position along the capillary is defined as the center of the dip in intensity seen in RBC shadows. The distance the RBC centers move along the capillary is determined manually by matching the dips for each pair of intensity traces. This distance is then used with the time between the image pairs (paired frame delay (PFD)) of the acquisition to calculate the velocity using equation 3

### F-Tilt Analysis

RBC angle was calculated in reference to the capillary for each frame pair using a custom MATLAB GUI. In the GUI, the user selects a small ROI containing the capillary of interest. In the select ROI, a reference line is drawn along the length of the capillary which defines the capillary angle α from equation 5. A second reference line is drawn along the edge of the RBC shadow which is used to define the angle *θ* in equation 5.

### Validation Experiment

F-Pair and F-Tilt estimates of RBC velocity were validated by comparing the estimated velocities from these two methods with the traditional line scan as demonstrated by Kleinfeld et al^12^. First, conventional raster scanning imaging is used to visualize the vasculature and identify an imaging region where capillaries are not obstructed by larger pial vessels. Capillaries with clear flow were identified, and the microscope was configured to perform a 15 μm line scan along the length of the selected capillary at 2.5 kHz for 1 second. This is then followed by two staggered scans with the first stagger scan having the fast axis oriented along the X axis and the second having fast axis oriented along the Y. This orthogonal scan patterns allows for the captures of flow in capillaries of any orientation. For both stagger scans, a line period of 512 ms (plus flyback overhead), line jump of 2, total line number of 256, and field of view size of 237.88 μm was used. For each capillary line scan, the acquisition was repeated for 2 minutes, giving 55 measurements per capillary. Raw intensity data was saved for post processing and analysis.

In post analysis the line scan data was reconstructed to provide a length vs time image of RBC shadows moving along the length of the capillary. Following an SVD based approach described previously^12^, the average RBC velocity over the 1 second scan is calculated. From the stagger scan images, RBC velocities were estimated using both F-pair and F-tilt methods for the same capillary that was measured with the line scan. These velocities were then compared with the line scan estimated velocities.

### Estimating Flow Speed Variability

To estimate flow speed variability, two minutes of stagger scans in 4 ROIs across 3 healthy adult mice were acquired. Similar to the validation experiment, ROIs were first identified using raster scanning before switching to stagger scanning. Capillaries with RBC shadows of sufficient contrast were identified, and F-pair and F-Tilt speed estimates were calculated giving speed traces for 40 capillaries across the fields of view.

### Capturing Capillary Stalling Parameters

To estimate capillary stalling parameters, ten-minute scans of a 713×713 μm at 0.57 Hz (512×512 resolution, 4 μs dwell time) were acquired and stalls were manually recorded as time points when RBCs in a capillary were stationary for consecutive frames. Similar to previous work using OCT^8,26^, we used a custom Matlab GUI to mark stalled capillaries and annotate all the time points where flow was stalled.

## Results

### Validation of Flow speed measurements

To quantify the flow speed accuracy obtained with F-Tilt and F-Pair from the stagger scanning method, a 1 second line scan over a select capillary is interleaved with the stagger scan acquisition sequence. From the line scan, the average flow speed is calculated^12,13^ and compared to velocities calculated using both the F-pair and F-Tilt from the stagger scans with a 237×237×120 μm volume and a paired frame delay of 5.56 ms. Figure 4 summarizes the results of the comparison across measurements taken from 24 different capillaries, with speeds ranging from 0-2 mm/s as measured using the line scanning technique. Figure 4b shows accurate F-pair velocity estimates in comparison to the line scanned estimates for capillaries with slower speeds, but F-Pair tends to underestimate velocities at higher speeds. This is because tracking movement using the frame pairs can underestimate higher speeds due to aliasing of the RBC shadows. Figure 4c compares the RBC velocity extracted using F-tilt against the velocity estimated by line scanning. Using F-Tilt, RBC velocities at higher speeds match velocities estimated by line scans and overestimates RBCs at slower speeds because shallower slew angles are harder to estimate from the image. Increasing the sampling resolution would give a better angle estimate at the expense of imaging speed. Therefore, we used of a combination of the two methods, taking the F-pair based estimate at slower speeds, and F-Tilt based estimate at faster speeds to better ensure accurate reported speeds (Figure 4d). This was done by adjusting the threshold for using the F-Tilt estimate vs. the F-pair estimate. If the estimated F-Tilt speed was above the cutoff, the F-Tilt speed was used. Below the cutoff, the F-pair speed was used instead. Using a cutoff of 0.4 mm/s was found to provide to minimize the error in speed estimate when using the line scan speed as ground truth (Figure 4d). We also found that accounting for the orientation of the capillary using Equation 6 is critical for an accurate F-Tilt estimation (Supplemental Figure 2).

**Figure 4.**
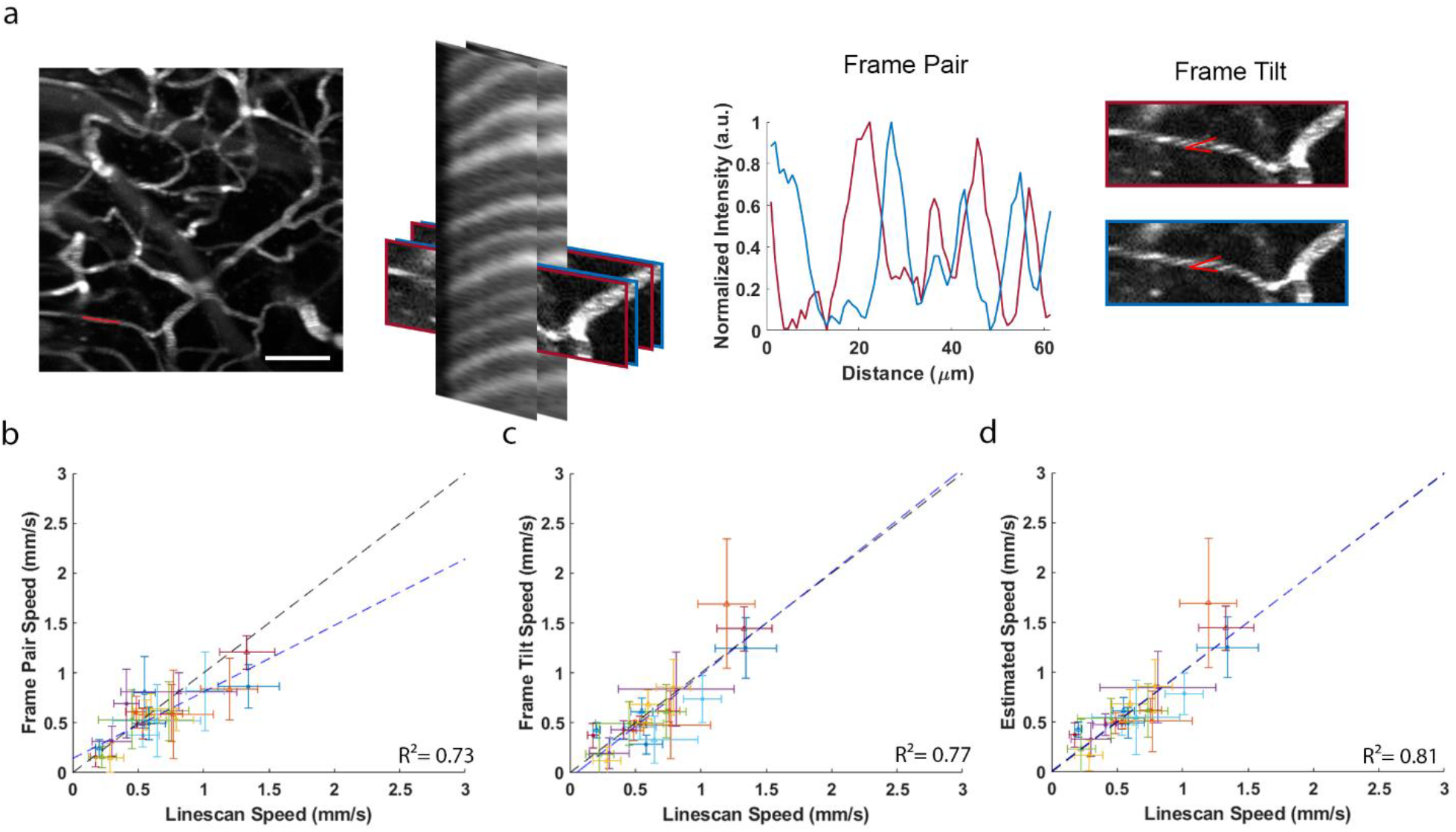
**(a)** Summary of the flow speed validation. For each sequence, the F-pair and F-Tilt measurement was compared to the line scan of a selected capillary interleaved between orthogonal stagger scans. **(b)** Comparison of the F-Pair speed with the line scan speed across capillaries. **(c)** Comparison of the F-Tilt speed with the line scan speed across capillaries **(d)** Line Scan Speed compared to a combined F-pair and F-Tilt estimate. For each time point, F-Tilt speeds > 0.4 mm/s were used, and F-pair speed used in all other cases.

### Tracking of capillary flow dynamics

Using a combination of F-pair and F-Tilt for estimating flow speeds, we track the flow dynamics within capillaries with a range of flow speeds (Figure 5 a-b), including in capillaries where flow undergoes abrupt changes when it stalls. Figure 5d shows an example time course of capillary flow speed estimated using our method compared to line scanning, including where a stoppage or “stall” and recovery of capillary flow occurred during the validation measurements. The similarity in reported flow speeds demonstrates that our approach can capture these dynamics when compared to a line scan taken of a single stalling capillary. We found the variability in flow measured in a capillary (measured as the coefficient of variation) to be well matched between line scans and stagger scans (Supplemental Figure 3), despite the fact that the line scans and stagger scans were performed sequentially and flow speed changes could occur when switching between the scans Figure 5 (e-h). However, in the case of staggered scanning, it is not limited to serially tracking single capillaries, as is the case with line scanning. Tracking the capillary network flow changes during these events would be of interest in the study of microvascular changes in both healthy animals and disease models.

**Figure 5.**
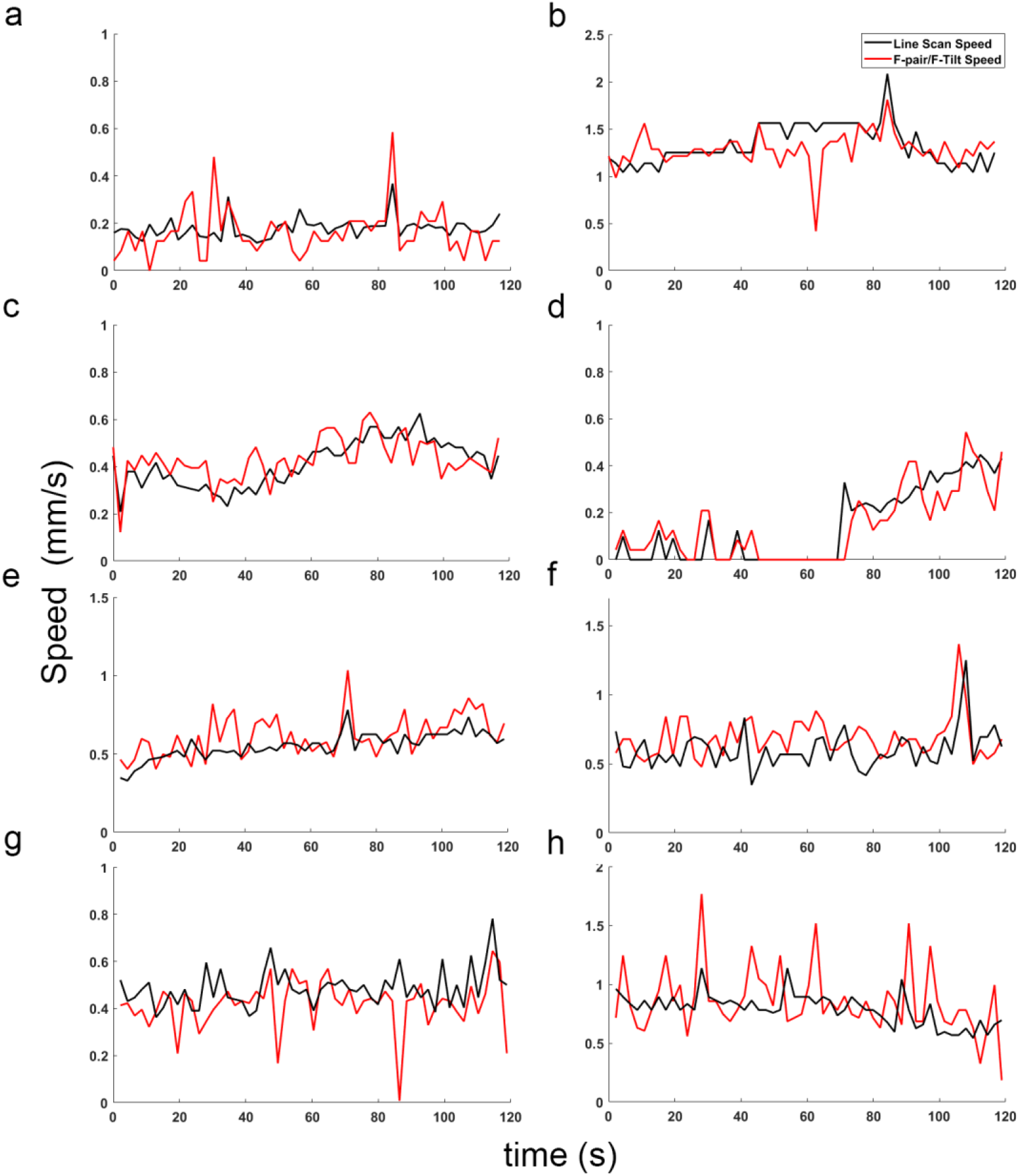
Tracking of Capillary Flow Dynamics. Comparison of the speed using the F-pair/F-Tilt estimate to the line scan speed averaged over 1 second in a **(a)** low speed capillary **(b)** high speed capillary **(c)** Capillary oriented at a 18.6° angle with respect to the fast axis of scanning and **(d)** Capillary whose flow stalled during the measurement **(e-f)** Capillaries where flow changes and variability are similar between stagger and line scans **(g-h)** Capillaries where fast changes in speed occurred between the stagger and line scans taken in sequence, though average speeds are similar.

### Tracking of capillary flow distributions

From modeling of capillary blood flow^9^, increases in transit time heterogeneity is associated with reduced oxygen delivery and has been observed in older mice where hypoxic pockets formed^5^. Using staggered scanning, we measured capillary flow speed in 4 fields of view between 3 healthy adult mice for 2 minutes per volume. Flow speed across an average of 10 capillaries per field of view was extracted and flow heterogeneity was estimated using the mean and standard deviation of flow both within and across the capillaries (40 capillaries in total). We found the mean flow speed across all capillaries to be 0.57 +/− 0.25 mm/s; an inter-capillary flow variability (the coefficient of variation of mean flow speed across capillaries) to be 0.42; and an intra-capillary variability (coefficient of variation of flow speed in a single capillary) to be 0.44+/− 0.2. Our measured inter capillary variability is similar to that reported using point scanning^11^ and line scanning^5^, which reported an inter-capillary (spatial) heterogeneity of ~38% and ~50% reported in young mice respectively. Interestingly, our measured intra capillary variability was significantly higher^11^ and indicates that the variability in flow speed within a given capillary is similar to the variability across the capillary network, at just under 50% of the mean flow speed. The implication is that across even a few minutes, capillary flow speed can vary significantly within a capillary, at a similar magnitude to the variability seen across the capillary network. The long, continuous measurement of capillaries with stagger scan allows us to observe these different capillary flow dynamics, since we do not need to change focus or measure capillaries one at a time.

### Estimation of capillary stalling parameters and effects

In addition to the capture of capillary flow speeds using staggered scanning, our Bessel beam based two-photon system provides an improved ability to catch stalling events compared to previously used techniques. Using standard raster scanning, we were able to image up to a 713×713×120 μm volume of the micro-vasculature at a rate of 0.57 Hz, generating angiograms where flow in >200 capillaries are visible on average (Figure 7a). This volume rate is a significant improvement over previously used OCT based angiograms (because of the shorter pixel dwell time), as well as standard two-photon microscopy which requires serially scanning in depth to acquire a volume. We used this large FOV to capture stalling events that occurred over a 10-minute time period (Figure 7b) and compared them to what has been reported previously using OCT-based methods^8,26^. Stalls were manually identified and classified as when an RBC shadow remained stationary for at least two sequential images (Figure 7c). Table 1 shows the results of the stalling analysis performed in 4 ROIs across 2 mice. As expected, we found an incidence comparable to that found with OCT based angiograms, and significantly larger than rates that have been observed with conventional two-photon microscopy^7,37^ since capillaries were monitored continuously for the full measurement time. The cumulative stall duration of an individual capillary was also found to be similar, though the point prevalence (average number of capillaries stalled at a given time) was found to be close to double. This is likely due to the higher frame rate of our imaging enabling us to identify shorter duration stalls than observable with the ~9 seconds per image with the OCT angiograms.

**Figure 6.**
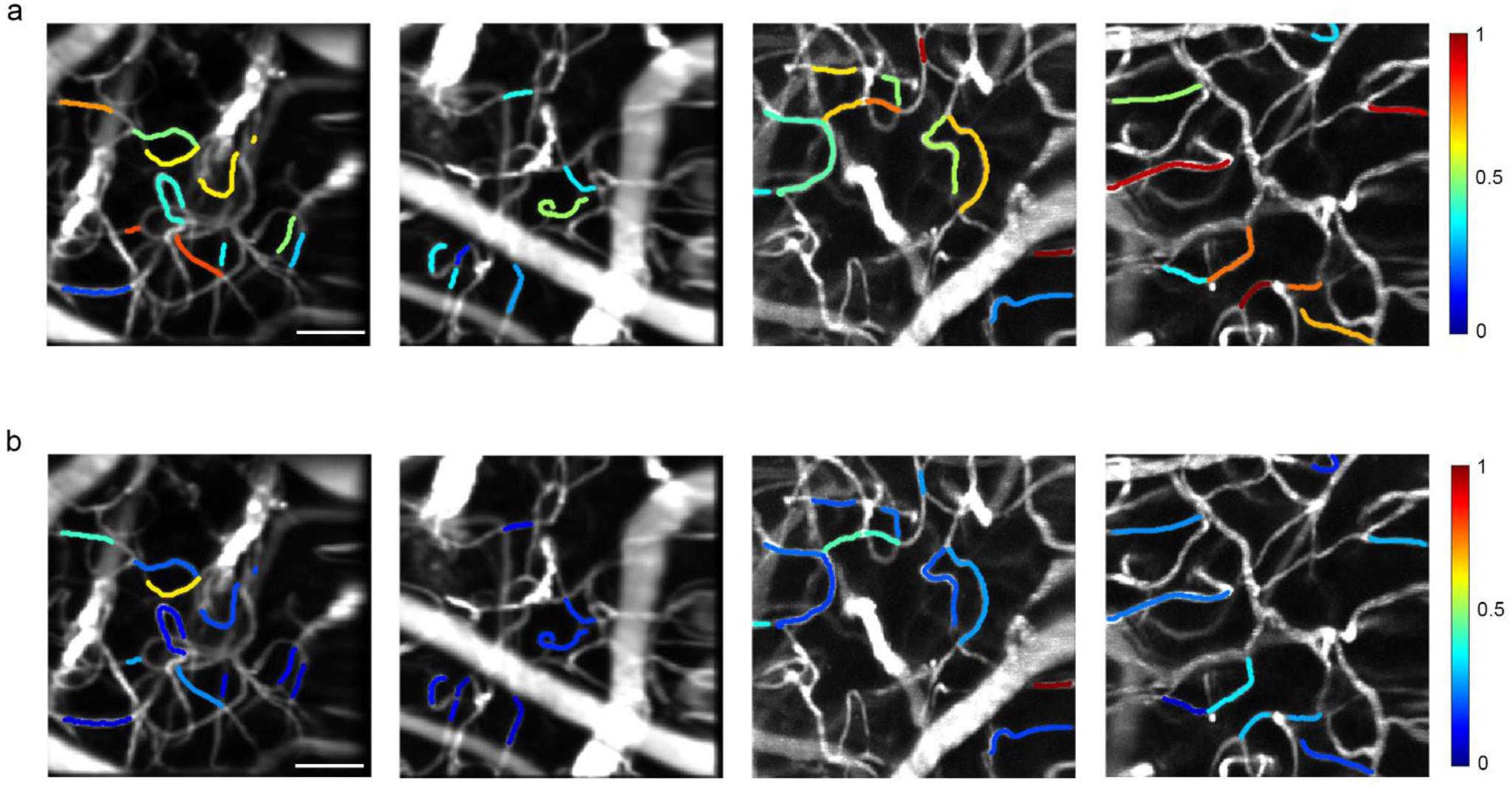
Comparison of flow speeds and flow variability in capillaries where F-pair and F-Tilt estimates could be obtained across 4 healthy adult mice **(a)** Mean capillary flow speed **(b)** Standard deviation of flow in the same capillaries across the 2-minute measurements

**Figure 7.**
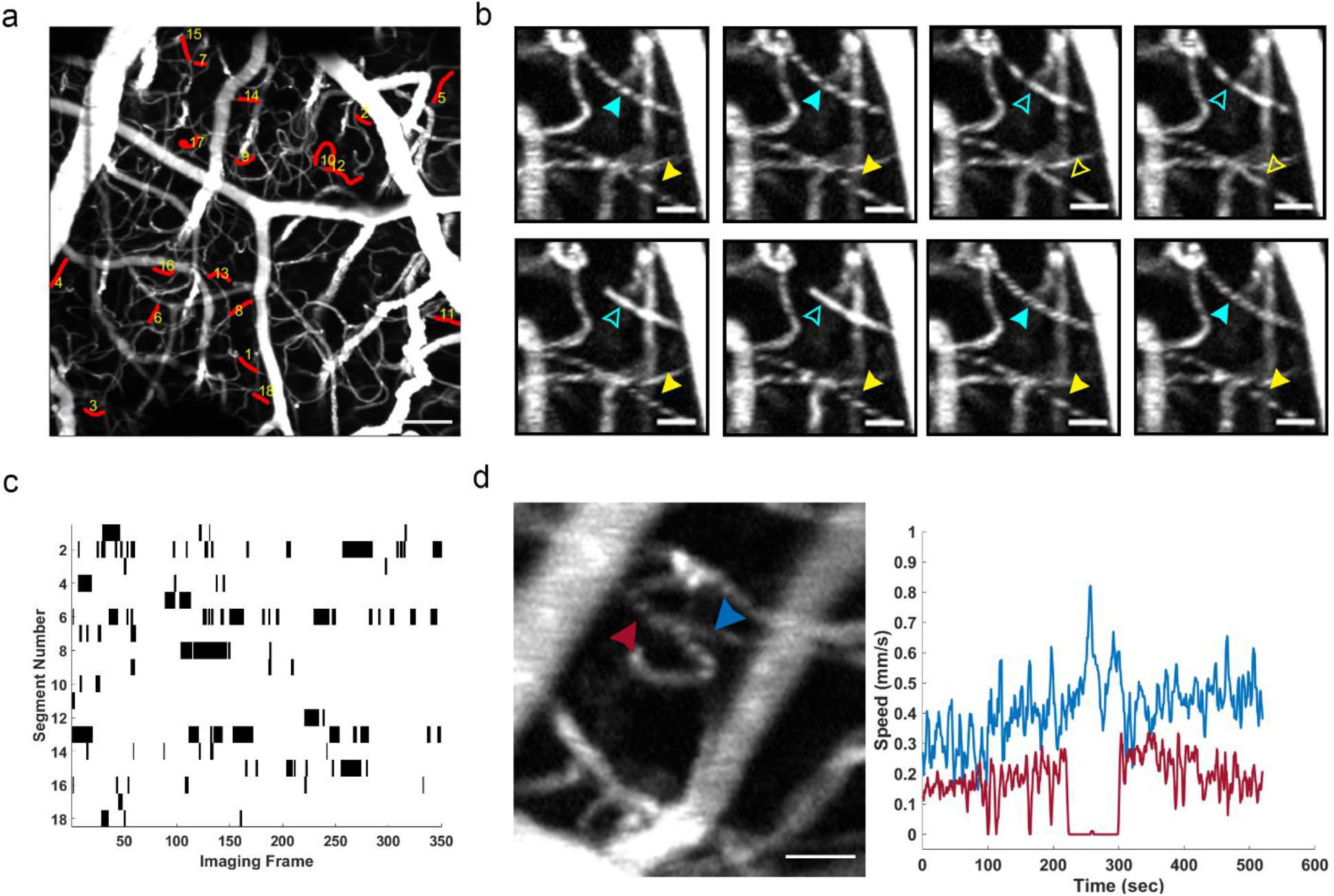
Capturing of capillary stalling event and flow dynamics **(a)** MIP of a 713×713×120 μm volume that was monitored using standard raster scanning for the identification of stalling events. Red highlighted segments represent capillaries that were identified to have stalled during the 10-minute measurement. **(b)** Example of two capillaries that experience stalling, indicated by the blue and yellow arrows. Solid arrows denote frames where the capillary is flowing, and hollow arrows denote times where RBCs are stalled in the capillary scale bar 20 μm **(c)** Stallogram showing the time points where each of the 18 identified stalls occurred. Black indicates time when flow was stalled. **(d)** Speed traces from a 10-minute stagger scan where a capillary stalled for longer than a minute, 4 frame running average. The connected capillary (blue arrow) flow speed increase during periods of slow/no flow in the stalling capillary (red). Pearson Correlation −0.31.

Previous work^7,8,26^ only quantified the occurrence rate of these events, but not their effect on nearby flow speeds. Using stagger scanning we were able to observe a change in flow speed in a capillary due to nearby stalling. Figure 7d shows an ROI when we capture a stalling of flow in a capillary that lasts for ~1 minute. During the time, we can see a significant increase in flow in a connected capillary during the stall, before returning to a lower speed when flow resumes in the stalled capillary. Finally, by using higher dwell times with our system, generating more fluorescence, we are better able to image capillary flow deep into the brain. Figure 8 shows a comparison of signal quality vs depth for single scans with a dwell time of 4 μs. RBC shadows are clear even after shifting the focus of the Bessel up to 350 μm below the surface. Below this (Figure 8d), capillaries are visible but the poor signal to noise means RBCs are not visible. We used F-Tilt to estimate speed at the maximum imaging depth (Figure 8d) in two representative capillaries (Figure 8d inset; Figure 8f) across ~ 1 minute of raster scanning, showing speed dynamics deep in the capillary network. However, the larger pixel size in these raster images leads to reduced sensitivity to small speed changes (Equations 4 and 5), and the lower slow axis line rate means a steeper slewing effect occurs at lower speeds, limiting the potential maximum speeds. Therefore while this demonstrates that stalling and flow speed data can be extracted across multiple cortical layers^13^ with our system, stagger scans with higher resolution would offer a more robust estimate of flow speed.

**Figure 8.**
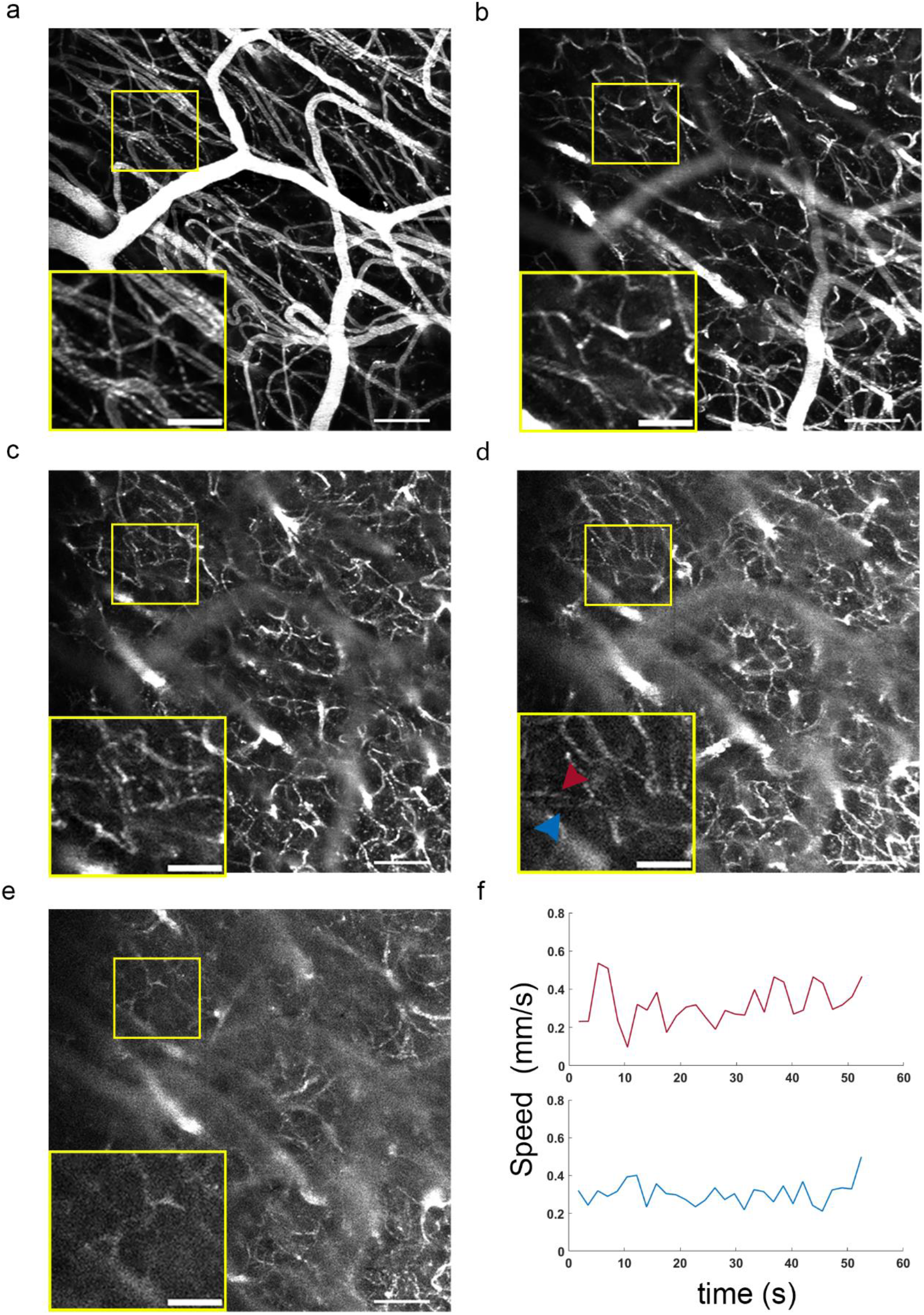
Resolving capillary flow across depth. Single 713×713×120 μm volume raster scanned with a 4 μs pixel dwell time at an approximate depth range of (**a**) 0-120 μm **(b)** 100-220 μm **(c)** 200-320 μm **(d**) 320-420 μm, and **(e)** 400-520 μm. Scale Bar 100 μm Inset: 50 μm (f) Speed measurement taken using F-Tilt for the capillaries indicated in the inset in (d).

|  | OCT (Erdener et al. 2017) | Bessel Beam TPM |
| --- | --- | --- |
| Number Animals | 6 | 2 |
| ROIs/Animal | 2 | 2 |
| Incidence | 6.8% +/- 2.9 | 9.94% +/- 1.35 |
| Point Prevalence | 0.31% +/- 0.7 | 1.15% +/- 0.38 |
| Cumulative Stall Duration | 10.6% +/- 3.2 | 11.7% +/- 4.37 |
| Mean Stall Duration | 23 sec +/- 15 | 9.39 sec +/- 17.12 |
| Frame Period | ~9 sec | 1.75 sec |

Comparison of stall parameters measured with OCT and Bessel Beam TPM

## Discussion

Previous studies of capillary flow dynamics, particularly around transient changes in flow, have been hindered by limited throughput or low frame rates. To overcome this, we built a custom two-photon Bessel beam microscope to enable efficient and reliable tracking of capillary flow speeds and capillary stalling events. Combined with our staggered scan pattern, we can track flow dynamics across the capillary network simultaneously at ~1 Hz. Further, the ability to perform additional fluorescent labeling opens the possibility for examination of the influence of other cell types on flow speed distribution^4,29,30^ using our technique.

The use of a staggered scan pattern to create frame pairs allowed us to capture and identify the flow of individual RBCs as they flow along a capillary while maintaining high dwell times to ensure a good signal to noise ratio. By tracking their displacement between each pair of frames, we can estimate RBC flow speed. Rather than being limited to scanning single capillaries or multiple capillaries in sequence, tracking can be done across multiple capillaries simultaneously. Using this technique, we show very reliable estimates of flow speed at speeds less than ~ 0.4 mm/s. Above this speed, this approach can underestimate speed in some cases, due to the aliasing of multiple red blood cell shadows that are present in the capillary. To help address this limitation, we use the angle of the RBC as it appears in the capillary to estimate its speed (F-Tilt). We found this better able to estimate faster flow speeds, but less accurate at lower flow speeds where the slewing effect was harder to distinguish. A combined approach was therefore used, resulting in reliable tracking over a large range of relevant flow speeds in capillaries across the field of view. With this combined approach, we were able to quantify flow speeds and similar levels of flow variability when compared to line scanning of individual capillaries (Supplemental Figure 3).

To leverage the ability for simultaneous flow speed measurements, we performed our scanning approach in 237×237×120 μm volumes and looked at flow variability across 40 capillaries in awake mice. Our results show capillary flow speed can vary significantly within a capillary over the course of 2 minutes, with a mean coefficient of variation (COV) of 0.44. We also found that the spatial variability (measured by the COV) to be very similar to the COV of mean speeds across the capillaries measured, and comparable to values reported with capillary line scans and point scan measurements^5,11^. Understanding changes in flow heterogeneity across the capillary network due to capillary stalling, vascular changes associated with aging^5^ and Alzheimer’s^38^, and vascular remodeling due to stroke^39,40^ will be key to mapping disease progression and the effectiveness of treatment. The ability to quantify both spatial and temporal variability in flow speeds simultaneously will provide important information on the health of the network. In addition, by increasing the number of capillaries where flow speeds can be measured, we also greatly increase the throughput for experiments. For the study of capillary stalling for example, stalling events occur randomly making line scanning ill-suited since it is limited to one or a few capillaries. This would require many sequential measurements in order to capture even a small number of these events. Instead, we can monitor a region of interest that contains many capillaries in parallel and analyze capillaries that stall during our measurements. Further, we can study the effects of stalling on capillaries nearby the stall (Figure 7). While this has been done with line scanning to study the effects of permanent occlusions^41,42^, the transient nature of capillary stalling requires the measurement of capillaries in parallel to capture similar dynamics.

We also demonstrated that our system is capable of monitoring hundreds of capillaries for the occurrence of stalling (Figure 7) using a large field of view conventional raster scanning pattern and identifying capillaries whose flow was disrupted for periods of time. These scans could then be followed up with stagger scans to measure flow dynamics. Since capillaries that stall have a high probability to stall again^26^, capillaries that stall during the large FOV survey scan are likely to stall again during a follow-up stagger scan measurement. Spatial and temporal flow variability can then also be quantified as a function of the stalling rate in a given region. This approach will greatly improve the ability to understand the impact of these stalls on flow distributions in the capillary network, as well as test the effects of functional activation and pharmaceutical intervention on capillary stalling and flow heterogeneity.

At our current line rate, we found the maximum measurable speed to be ~ 2 mm/s, making the majority of capillary flow speeds measurable^11,12^. Above this speed, we typically are unable to resolve individual RBCs in the capillary, limiting the ability to make a speed measurement using either F-pair or F-Tilt. This limitation could be overcome by using higher line rates, decreasing the time taken to scan over the capillary as the RBC flows through. However, the use of higher line rate always comes with the tradeoff of shorter dwell times and worse signal to noise ratio (SNR). Further, while we were able to use standard galvometers for our study, a significant increase in line rates would require the use of resonant scanners, removing the ability to rotate the scan field to align the fast axis with capillaries of arbitrary orientation. The capillaries that could be analyzed using our technique would then be limited based on their orientation to the resonant galvo axis. One needs to consider their desired measurement range, as well as signal to noise limitations when deciding on scan parameters. Alternatively, F-Tilt analysis could also be performed on conventional raster scans (Figure 8 f), including previously acquired data in any raster scanning system where the line rate is known. Compared to previous studies using a similar approach, we also demonstrated the ability to track much faster capillary flow speeds and have a higher frame rate^15,16^.

The scan pattern and analysis of RBC displacement (F-pair) and slewing (F-Tilt) due to flow during scanning are straightforward to implement and can be done with any scanning-based system, making it readily applicable for anyone interested in studying capillary flow or other faster dynamics where conventional frame rates are not sufficient. For example, three photon microscopy is limited due to need to use low repetition rate sources (typically 1-2 MHz), capping the maximum line rate of scanning^43–45^. The line rates used in our studies are compatible in this case and could easily enable high throughput studies of capillary flow speeds in deep brain regions. This is of particular importance as microvascular structure and flow dynamics can differ across brain regions^11,46^, as well as changes associated with aging^6^ and pathology^47^. The only additional technical requirement is the ability to control the scan path, making it easily adapted to any scanning based system. It also provides flexibility in the image resolution and pixel dwell time, making it tunable from experiment to experiment based on the field of view, dwell time, and temporal resolution required. In summary, we developed a scanning approach with a custom Bessel beam two-photon microscope that allows two different methods for robust tracking of flow speed across numerous capillaries simultaneously. We then applied these techniques and quantified both spatial and temporal flow variability in awake mice, as well as demonstrated the utility of the Bessel beam microscope in capturing stalls in capillary flow.

## Data and Code Availability

The data sets as well as the custom MATLAB code used to acquire, process, and analyze the data are available from the corresponding author on request.

## Acknowledgements

We thank Anna Devor her helpful discussion

## Author Contributions

JG, SP and AC equally contributed to the design and construction of the 2P Bessel beam microscope. JG, AC and DAB developed the stagger scan technique and the F-tilt/F-Pair models for estimating RBC velocity. JG wrote the Matlab code that adds stagger scanning to Scan Image and worked with SK to write the analysis code for extracting RBC velocities and stalls. JG acquired and analyzed all data with input from SK, DAB, and AC. KK and JJ performed all surgical procedures and provided animal assistance during experiments. JG, AC, and DAB wrote and edited the manuscript.

## Supplemental Figures

**Supplemental Figure 1.**
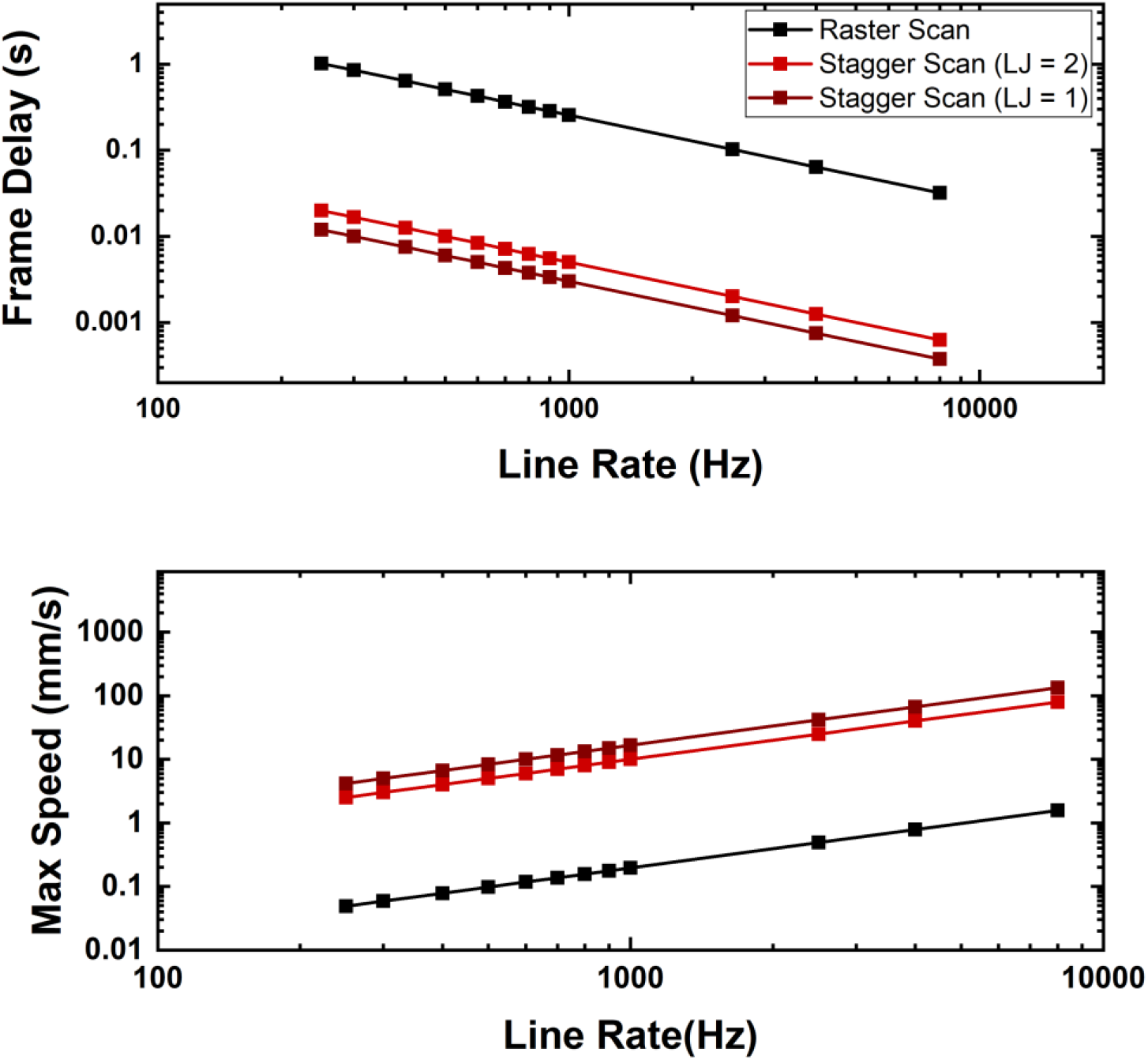
**(a)** Comparison of the time between frames in raster scanning and frame pairs in stagger scanning, based on the line rate of the scan. For the same line rate, stagger scans with line jumps of 2 (maroon) and 3(red) have significantly shorter time delays, calculated using Equation 1. **(b)** Comparison of the theoretical maximum flow speed measurable using raster scanning and Stagger scanning with line jumps of 2 and 3 lines. Max speed is based on a theoretical capillary that is 50 μm long, and a uniquely identifiable RBC flow from one end of the capillary to the other between a given pair of frames

**Supplemental Figure 2.**
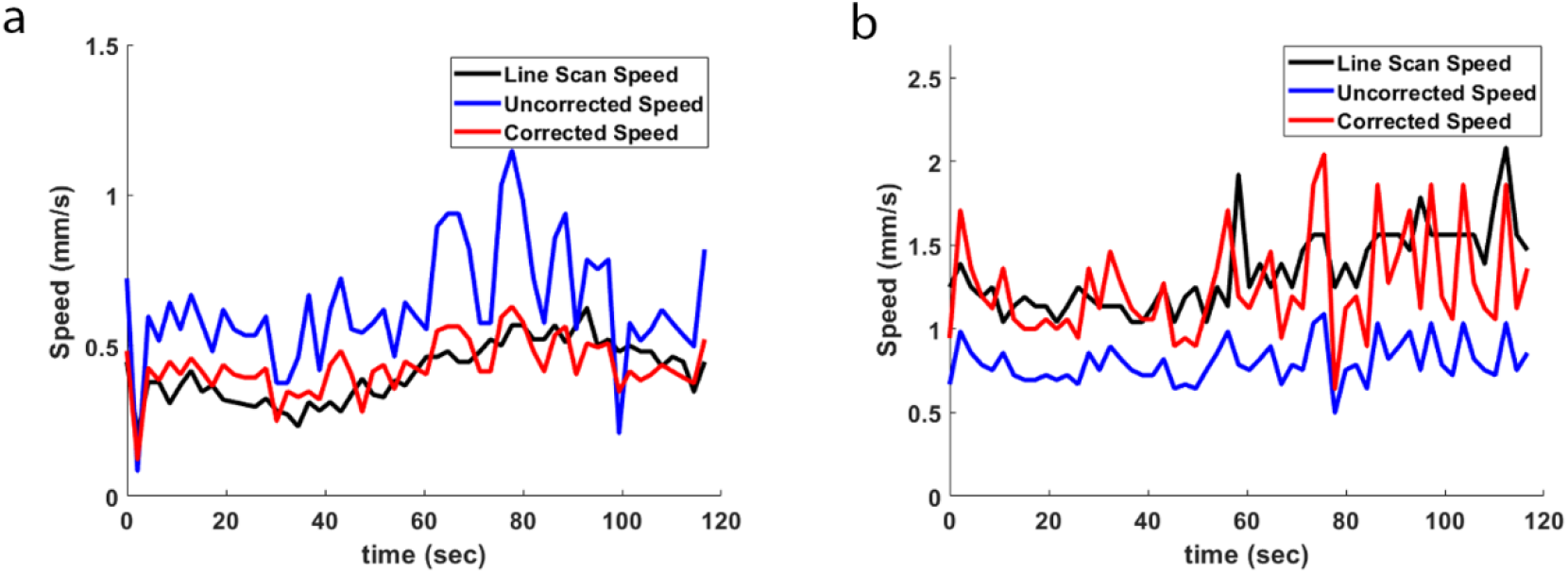
Comparison of flow estimates using the uncorrected and corrected F-Tilt estimate in a capillary with flow: **(a)** in the same direction as the slow axis scan direction, resulting in an overestimation of speed; **(b)** in the opposite direction of the slow axis of scanning, resulting in and underestimate of speed without correction.

**Supplemental Figure 3.**
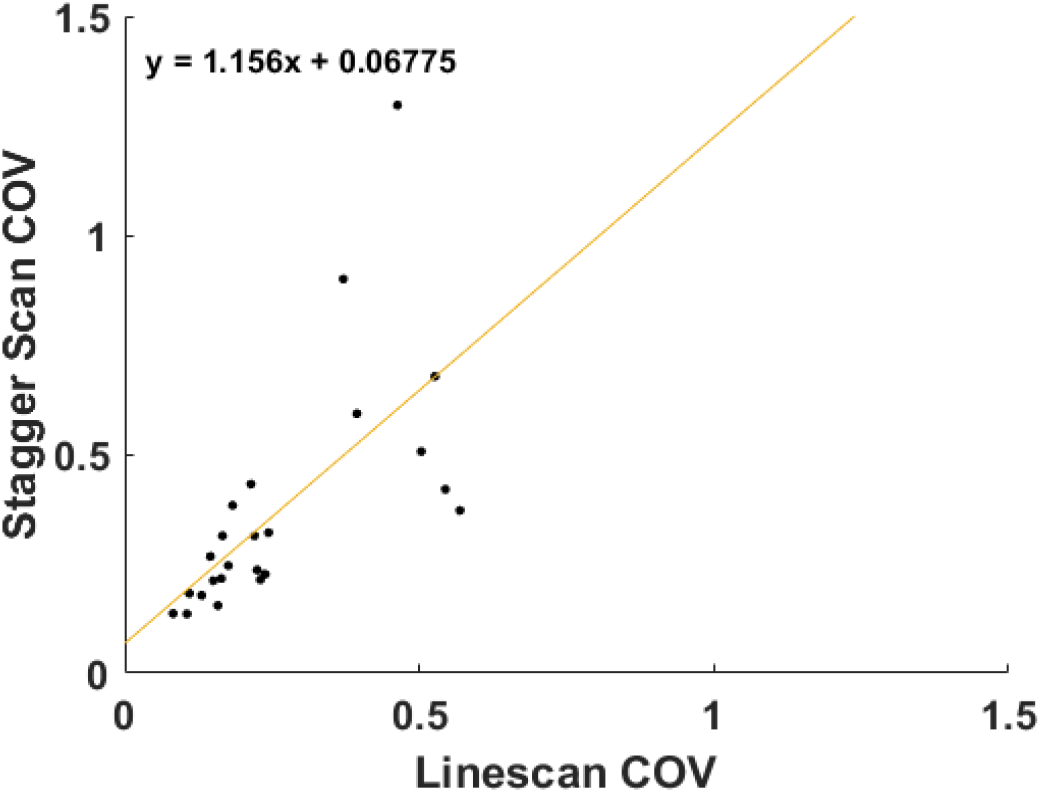
Comparison of the coefficient of variation calculated from stagger scan estimate vs line scan estimate in capillaries from the validation experiments.

